# CryoEM structure of the di-domain core of *Mycobacterium tuberculosis* polyketide synthase 13, essential for mycobacterial mycolic acid synthesis

**DOI:** 10.1101/2024.04.28.591520

**Authors:** Hannah E Johnston, Sarah M Batt, Alistair K Brown, Christos G Savva, Gurdyal S Besra, Klaus Fütterer

**Affiliations:** School of Biosciences and Institute of Microbiology and Infection, University of Birmingham, Birmingham, B15 2TT, UK; Biosciences Institute, Faculty of Medical Sciences, Newcastle upon Tyne, NE2 4HH, UK; Institute of Structural and Chemical Biology, The University of Leicester, University Road, Leicester, LE1 7RH, United Kingdom

## Abstract

Mycobacteria are known for their complex cell wall, which comprises layers of peptidoglycan, polysaccharides and unusual fatty acids known as mycolic acids that form their unique outer membrane. Polyketide synthase 13 (Pks13) of *Mycobacterium tuberculosis,* the bacterial organism causing tuberculosis (TB), catalyses the last step of mycolic acid synthesis prior to export to and assembly in the cell wall. Due to its essentiality, Pks13 is a target for several novel anti-tubercular inhibitors, but its 3D structure and catalytic reaction mechanism remain to be fully elucidated. Here, we report the molecular structure of the catalytic core domains of *M. tuberculosis* Pks13 (Mt-Pks13), determined by transmission cryo-electron microscopy (cryoEM) to a resolution of 3.4 Å. We observe a homodimeric assembly comprising the ketoacyl synthase (KS) domain at the centre, mediating dimerization, and the acyltransferase (AT) domains protruding in opposite directions from the central KS domain dimer. In addition to the KS-AT di-domains, the cryoEM map includes features not covered by the di-domain structural model that we predict to contain a dimeric domain with similarity to dehydratases, yet likely lacking catalytic function. Analytical ultracentrifugation data indicate a pH-dependent equilibrium between monomeric and dimeric assembly states, while comparison with the previously determined structures of *M. smegmatis* Pks13 indicates architectural flexibility. Combining the experimentally determined structure with modelling in AlphaFold2 suggests a structural scaffold with a relatively stable dimeric core, that combines with considerable conformational flexibility to facilitate the successive steps of the Claisen-type condensation reaction catalysed by Pks13.

## Introduction

Tuberculosis (TB) is caused by infection with *Mycobacterium tuberculosis* (*Mtb*) and continues to be a global health problem, prevalent in developing countries, and affecting millions of individuals every year. WHO-led efforts have gradually reduced the number of patients with active disease to about 10.6 million a year and the number of TB-linked deaths to about 1.3 million (WHO, 2023). However, infections with *Mtb* strains resistant to the standard treatment regimen remain high, accounting for about 3.3% of new and ∼17% of previously treated infections (WHO, 2023). It is widely assumed that developing novel antibiotics in combination with directing them at new cellular targets will be key to overcoming resistance, controlling the pandemic, and eventually eliminating TB. Novel therapeutic options arise from phenotypic screening campaigns, which probe growth inhibition and require subsequent target identification of inhibitory compounds. This approach has frequently returned proteins and enzymes involved in cell wall synthesis as candidates, because cell wall integrity is essential for pathogen survival in the infection host. Among these recently identified targets is polyketide synthase 13 (Pks13, Rv3800c), a multi-catalytic enzyme involved in the final stages of mycolic acid synthesis (Portevin et al., 2004).

Mycolic acids consist of a long **□**-hydroxy chain with a shorter **□**-alkyl side chain, in total containing between 60 to 90 carbon atoms, and form the outer membrane in the unique multi-layered cell wall of *Mtb* (Batt, Burke, et al., 2020). Analogous to the outer membrane of Gram-negative organisms, the ‘myco-membrane’ contains an ‘inner leaflet’ of mycolic acids that are covalently bound to and layered on top of arabinogalactan polymer, while the ‘outer leaflet’ is made of trehalose- or glycerol-esterified mycolic acids, interspersed with solvent-extractable free lipids (Batt, Burke, et al., 2020). Highly hydrophobic in character, the mycobacterial outer membrane provides an efficient barrier to antibiotics and the host defences, but pathogen survival is compromised when mycolic acid synthesis is suppressed (Brennan & Nikaido, 1995; Christensen et al., 1999; McNeil et al., 1991; Portevin et al., 2004).

Mycolic acids synthesis occurs in the bacterial cytosol, followed by export to the cell wall through the RND-family transport major membrane protein large 3 (MmpL3) in the form of trehalose monomycolate (TMM) (Grzegorzewicz et al., 2012; Varela et al., 2012). The synthetic pathway involves fatty acid synthase I (FAS-I) producing the short α-alkyl branch (C24-26) and fatty acid synthase II (FAS-II) synthesising the longer meromycolate chain (C56), which are joined together in the Claisen-type condensation reaction that is catalysed by Pks13 (Portevin et al., 2004).

Pks13 is an iterative type I polyketide synthase (PKS) composed of a sequence of catalytic and acyl carrier protein (ACP) domains separated by non-globular linker regions (Fig. 1A). The domain composition includes two functional ACPs, one located at the N-terminus and the other towards the C-terminus of the protein, while a non-functional ACP is located immediately upstream of the C-terminal thioesterase (TE) domain. Sandwiched between the functional ACPs are the central ketoacyl synthase (KS) and acyltransferase (AT) domains, which facilitate the condensation reaction (Portevin et al., 2004), while the TE domain at the C-terminus loads the newly formed mycolate onto a trehalose.

**Figure 1.**
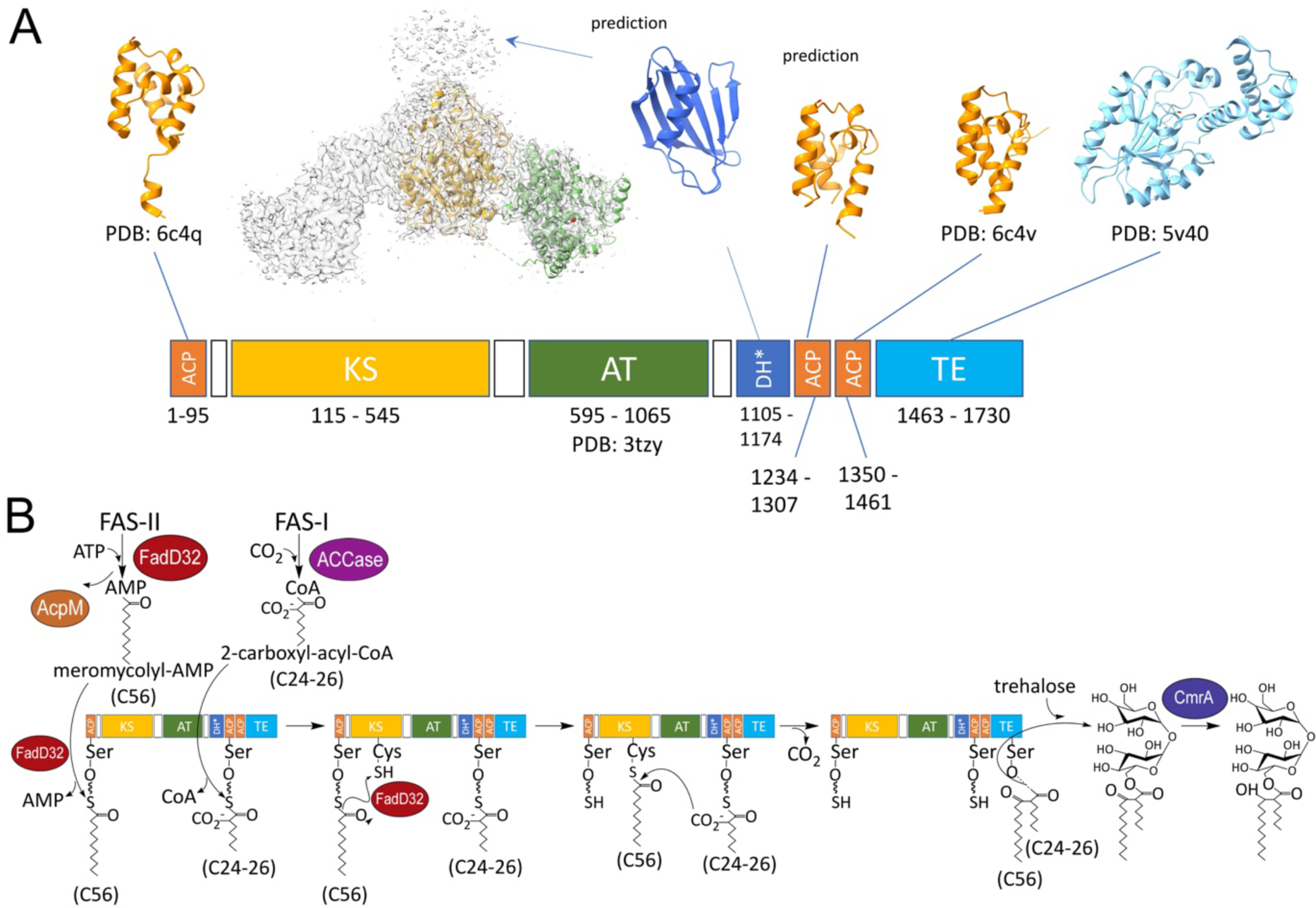
Overview of Pks13 domains, structure and reaction mechanism. **A** Domain architecture of Mt-Pks13 with experimentally derived or predicted structures. Abbreviations: ACP, acyl carrier protein; KS, ketoacyl synthase; AT, acyltransferase; DH*, pseudo-dehydratase; TE, thioesterase; ACCase, acyl-CoA carboxylase. The cryoEM structure from this study covers the KS-AT di-domain, a single monomer is shown within the recorded density of the dimer. The AlphaFold structure of the DH* fragment, is predicted to form contribute to dimer formation, consistent with the low resolution cryoEM map. The two ACPs and TE domain are crystal structures (PDB entries indicated) and the central ACP domain is an AlphaFold predicted model. **B** The proposed Pks13 catalytic mechanism. See text for details (phosphopantetheine, Ppant, is denoted by thick wavy line). Schematic adapted from (Batt, Minnikin, et al., 2020).

The proposed catalytic mechanism for Pks13 (Fig. 1B) commences when the phosphopantheine (Ppant) arms of the functional ACPs are loaded with the respective acyl chains. The meromycolate chain (C56) from FAS-II is activated with AMP and loaded onto the first ACP (Ser 55) by FadD32 (Léger et al., 2009) and then transferred to the active-site cysteine (Cys 287) of the KS domain, while the shorter α-alkyl chain (C24-26) from FAS-I is carboxylated and transferred to the second ACP (Ser 1266) in a step facilitated by the AT domain. The Claisen condensation reaction proceeds via a nucleophilic attack of the carbonyl group of the C56 chain by the acidic α-carbon of the C24-26 chain. Finally, the TE domain functions both as a hydrolase and an acyltransferase, cleaving the newly formed α-alkyl β-ketothioester intermediate from the C-terminal ACP and loading it onto trehalose (Gavalda et al., 2014). While the sequence of biochemical steps is known, how they are orchestrated by the multi-domain structure of Pks13 is unclear. Here, we report the cryoEM structure of the KS-AT di-domain of Mt-Pks13 to help clarify the molecular mechanism of the final step of mycolic acid synthesis.

## Results

### Pks13 purification required a three-step chromatography procedure

Mt-Pks13 was overexpressed in *E. coli* BL21 cells and purified in a three-step column chromatography process (Fig. 2). First, a nickel column, based on affinity of a hexa-histidine tag on the C-terminus of the protein, separated Mt-Pks13 from the bacterial cleared lysate. A QHP ion-exchange column, further purified the protein (Fig. 2B). In a subsequent size exclusion chromatography (SEC) step, Mt-Pks13 eluted as a single peak (Fig. 2C) of approximate 95% purity based on SDS-polyacrylamide gel electrophoresis (SDS-PAGE) of the peak fractions (Fig. 2C). Presence or absence of reducing agent had a profound effect on the quality and solution behaviour of Pks13. Initially, in the absence of a reducing agent, Pks13 Mt-Pks13 eluted in the void volume (∼104 mL, Fig. 2D), whereas addition of 2.5 mM β-mercaptoethanol to the purification buffers led to a shift of the elution peak for Mt-Pks13 to 125 mL, and resulted in higher purity as judged by SDS-PAGE.

**Figure 2.**
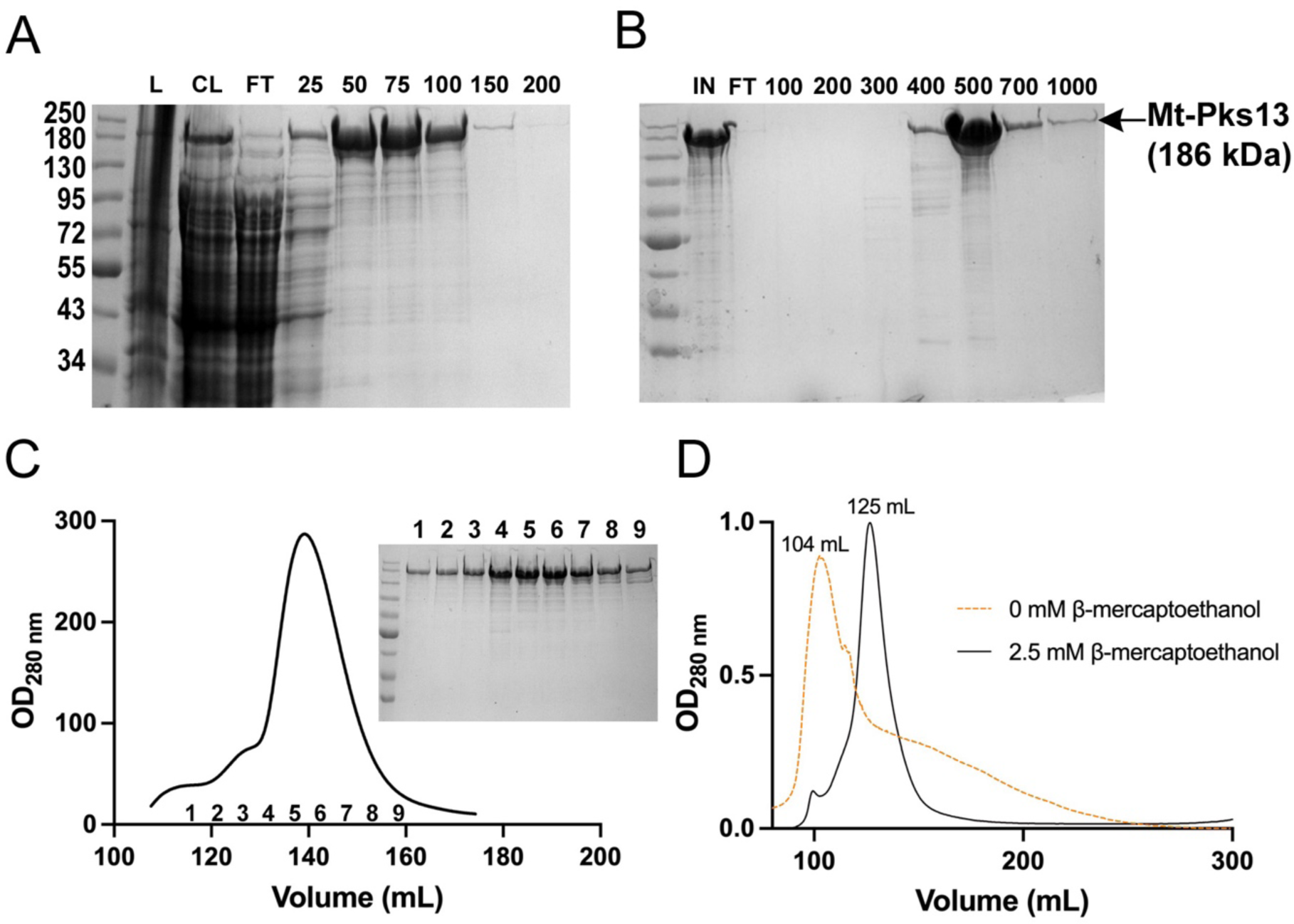
Three-step column purification of Pks13. **A** Ni-NTA column. SDS-PAGE of elutions from a nickel affinity column (L, lysis and CL, clarified lysate from *E. coli*; FT, flow through). The imidazole gradient (25-200 mM) is labelled above. **B** QHP ion-exchange column. SDS-PAGE of the elution samples from the column (IN represents the sample loaded onto the column) and the NaCl elution gradient is labelled above (100-1000 mM). **C** Size exclusion column (SEC) using a Superdex 200 column. Absorbance measured at 280 nm shows a single elution peak between 120-160 mL. SDS-PAGE of the corresponding fractions confirms pure Mt-Pks13. NEB Prestained protein standard (New England Biolabs) was used for all gels, while Mt-Pks13 (186 kDa) is labelled. **D** Chromatogram on a Sephacryl S300 HR (16/60) size exclusion column, measuring absorbance at 280 nm, illustrating the effect of β-mercaptoethanol on the elution behaviour of Mt-Pks13.

### Mt-Pks13 shows pH-dependent self-association *in vitro*

Analytical ultracentrifugation (AUC) in sedimentation velocity mode showed a systematic and substantial underestimate of the sequence-derived molecular mass (Mr) of Mt-Pks13 of 63.5 kDa at pH 7.9 (Fig. 3). Since molecular shape considerably affects mass estimates derived from velocity data, we conducted an analysis in sedimentation equilibrium mode to eliminate the shape factor (Fig. 3A, 3B), where the protein was kept in its purification buffer (pH 7.9). In this case, the experimentally determined mass of 187.6 ± 2.5 kDa closely matched the sequence-derived mass for the Mt-Pks13 monomer of 186.5 kDa, with the recombinant poly-histidine affinity tag accounting for the small discrepancy between the two values. The close match between experimentally determined and sequence-derived mass also indicated that the massive mass underestimate in the velocity data are not likely the result of protein degradation during the AUC experiment.

**Figure 3.**
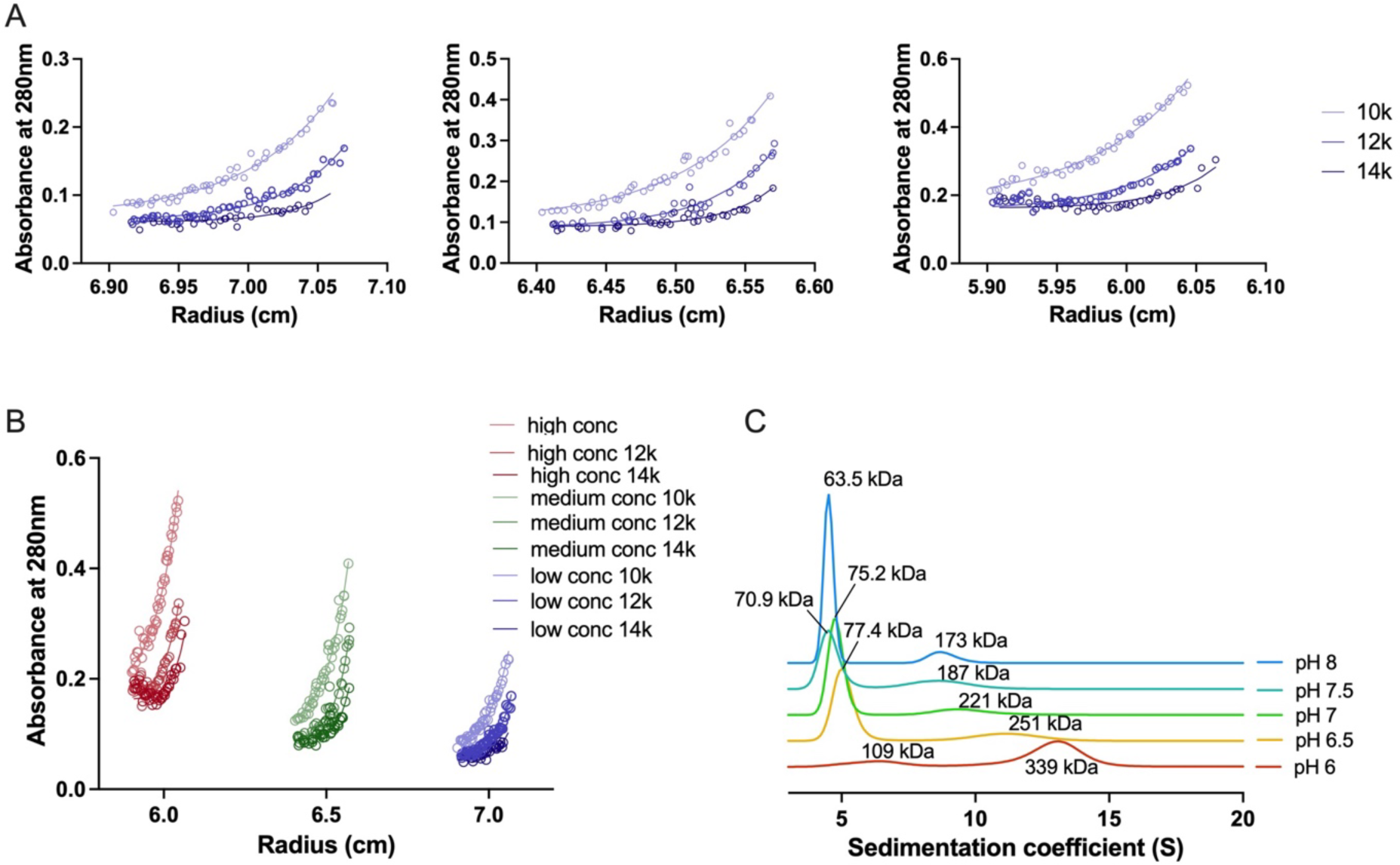
Analytical ultracentrifugation of Mt-Pks in sedimentation equilibrium and velocity modes. **A, B** Sedimentation equilibrium experiment of Mt-Pks13. The experiment was performed at pH 7.9 for low (A_280_ 0.25), medium (A_280_ 0.5) and high (A_280_ 0.75) protein concentrations, using three centrifugal forces (10, 12 and 14k rpm) at 4°C. **B** is a compilation of the sedimentation equilibrium results. **C** Sedimentation velocity experiment of Mt-Pks13, varying the pH between 6 and 8.

Nonetheless, the monomeric assembly apparent from the AUC equilibrium data stood in contrast to dimeric assembly states of PKS-type enzyme described in the literature (Tang et al, 2007). We wondered whether pH might affect the assembly state and examined self-association as a function of pH (Fig. 3C). The sedimentation velocity traces recorded over a pH range from pH 6 to 8 show a clearly discernible progression from a dominant high molecular weight peak (∼330 kDa at pH 6) to a dominant low molecular weight peak (∼100 kDa at pH 6) as pH becomes more alkaline. We interpret this finding as a dimer to monomer transition as pH changes from acidic to alkaline, considering the shape effect on mass estimates and the sedimentation equilibrium data.

### CryoEM structure of the KS-AT domain of Pks13

Samples of flash frozen full-length Mt-Pks13 expressed in *E. coli*, and analysed by single-particle cryoEM led to a 3D reconstruction of the KS-AT di-domain. Around 1400 electron micrographs were recorded from which several rounds of particle picking resulted in a data set encompassing ∼168,000 particles (Supplementary Fig. 1) that was used to calculate the final density map with an overall resolution of 3.4 Å.

### Overview of the Mt-Pks13 cryoEM map

The electron density map derived from 3D reconstructions of the electron micrographs shows a dimeric core domain with approximate 2-fold rotational symmetry (Fig. 4A), where the 2-fold rotation axis is oriented vertically in the top panel of Fig. 4A, and directed towards the viewer in the bottom panel. Indeed, two-fold symmetric projections readily emerged from the 2D class averages (Supplementary Fig. 1) without applying symmetry constraints in particle averaging, suggesting a considerable if not dominant population of dimeric species among particles captured on the cryoEM grid.

**Figure 4.**
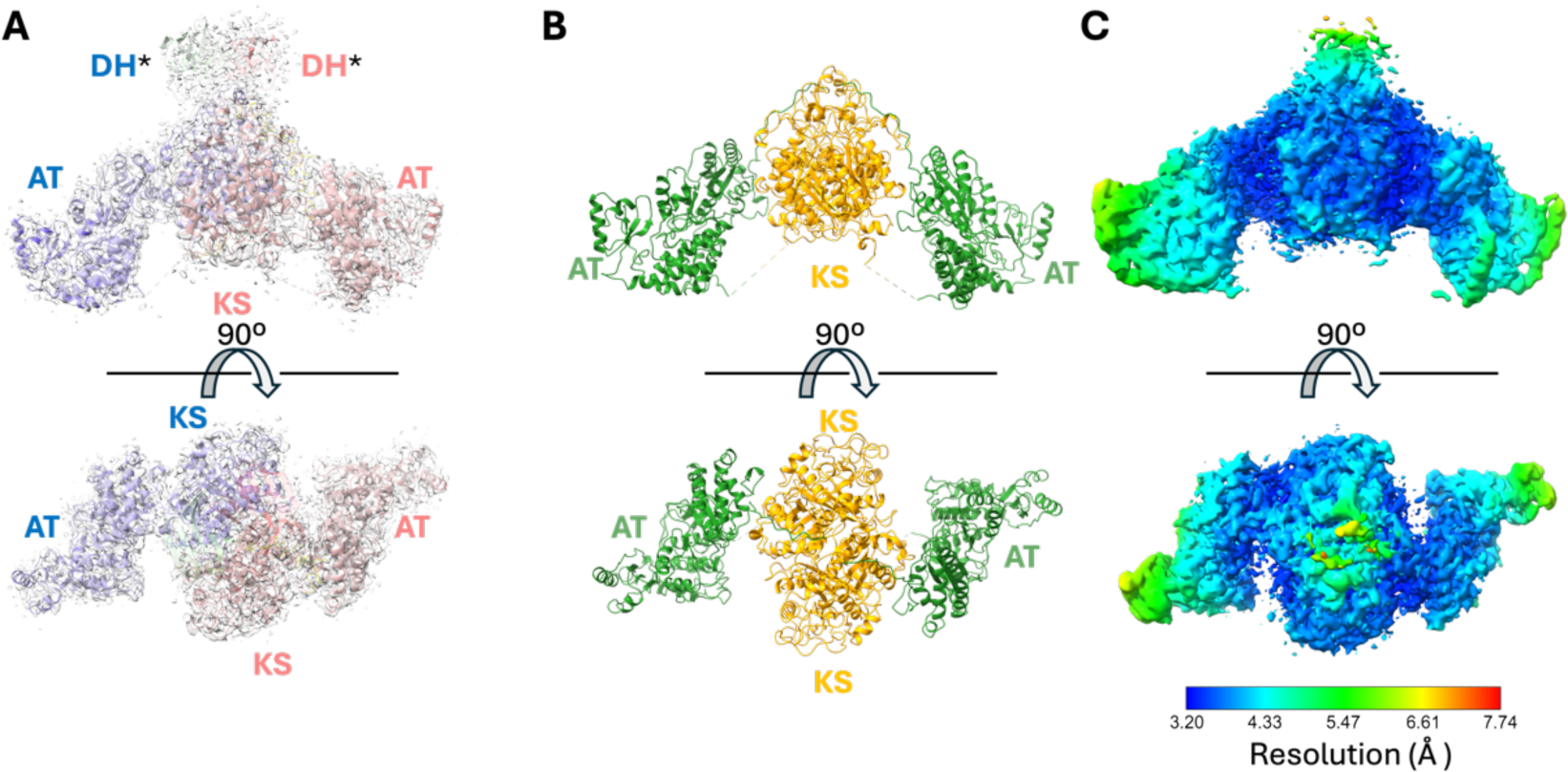
Electron density map of the dimeric di-domain assembly of Mt-Pks13 and analysis of local spatial resolution. **A** High resolution density map of the KS-AT di-domain, with ribbons coloured by chain. **B** Ribbon diagram of the KS-AT di-domain: KS (yellow) and AT (green). **C** Corresponding space-filling model of the structure, coloured by resolution (key at the bottom). Views from above and the side. Abbreviations: KS, ketoacyl synthase; AT, acyltransferase; DH*, pseudo-dehydratase domain.

The overall resolution of the electron density map derived from 3D reconstruction was about 3.4 Å, although the local resolution varied considerable across the observed dimeric assembly (Fig. 4C). The central KS domain dimer is resolved best, with local resolution down to 3.2 Å, while the distal ends of the AT domains show a spatial resolution of around 5 Å.

The dimeric core encompasses the KS and AT domains, which are sufficiently defined to build a structural model and assign the amino acid sequence to the protein backbone. The structural model covers residues 106-550, and 596-1073 of the amino acid sequence of Mt-Pks13 for both monomers. Dimerisation of the structural core of Pks13 is mediated by the central KS domain, flanked on either side by the AT domains (Fig. 4A, 4B). Above the KS dimer (in the orientation of Fig. 4A) we find distinct yet ‘untraceable’ density that is not covered by either of the two core domains.

Structure predictions using AlphaFold2 attribute this density to a structural domain that bears similarity to dehydratase domains in other polyketide synthases. We designate this domain as the DH* domain (pseudo-dehydratase). In the high-resolution map, density unequivocally linking the KS to the AT domain is missing for both monomers, but the sequence linking the AT to the DH* is traceable, including side chain orientations (Fig. 5).

**Figure 5.**
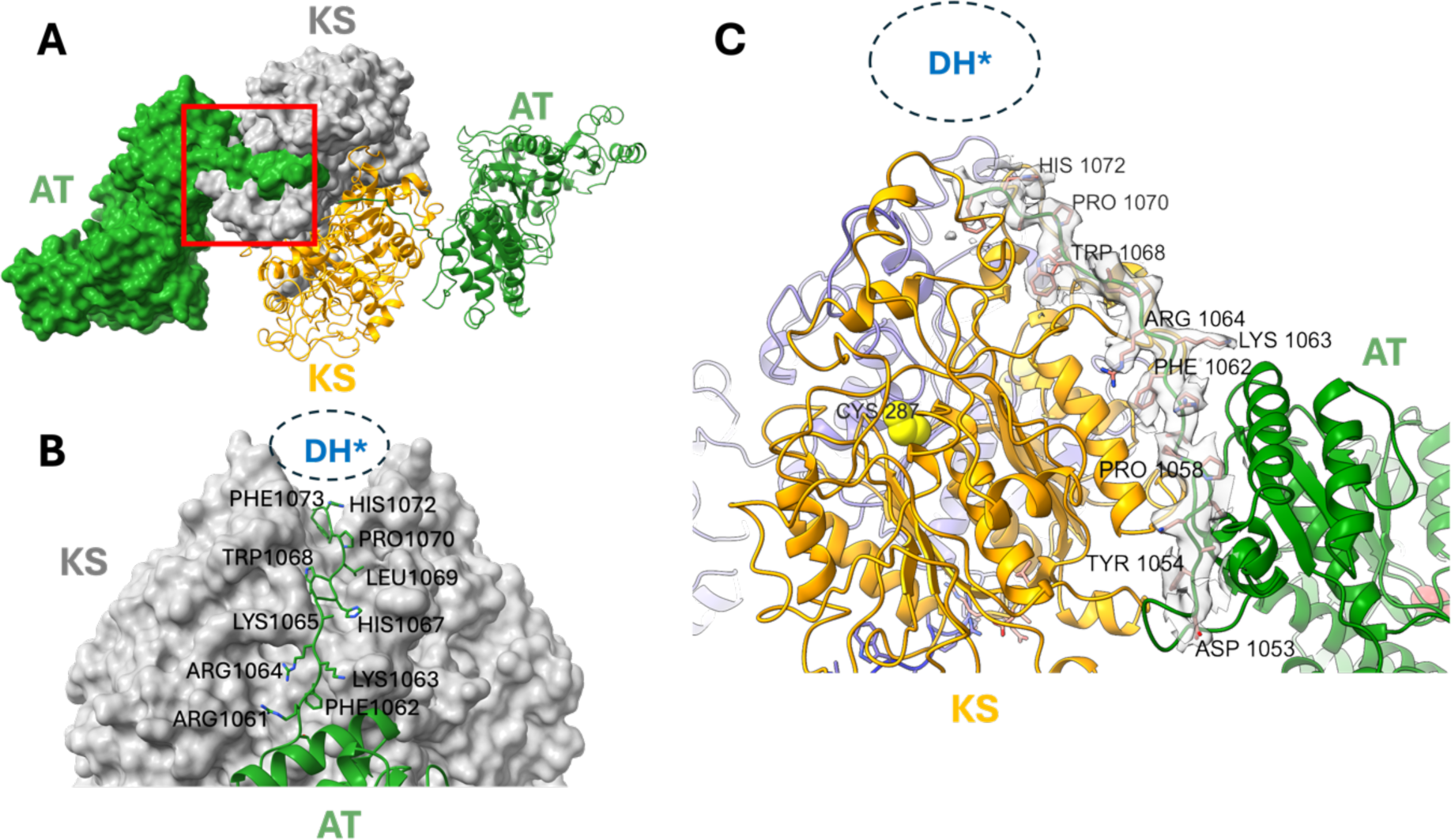
Linker between the AT and DH* domains. **A** View down the 2-fold symmetry axis of the Mt-Pks13 dimer in a mixed surface and ribbon representation, illustrating the position of residues 1053-1073 (the AT-DH* linker) in relation to the AT and KS domains. **B** Close-up view of the AT-DH* linker, representing the red boxed area in panel A, with selected amino acids labelled. **C** CryoEM density map for the AT-DH* linker. Abbreviations: KS, ketoacyl synthase; AT, acyl transferase; DH* pseudo-dehydratase.

### Dimeric di-domain KS-AT

The present structure indicates that dimerisation of Pks13 is mediated chiefly by the KS domain, while the DH* domain may also contribute to the dimer interface. Analysis of the KS dimer interface using PISA (Krissinel & Henrick, 2007) indicates a buried surface area of around 3500 Å^2^ per monomer. Despite the substantial burial of solvent-exposed surface area upon complex formation, the Complex Formation Significance Score (CSS) calculated by PISA is 0, suggesting that dimerization may not be constitutive, consistent with the pH-dependent monomer-dimer equilibrium observed in the sedimentation velocity experiments (Fig. 3). Reversible self-association aligns with a largely polar character of the dimer interface (Fig. 6A), which includes a small set of salt bridge interactions and about 35 H-bonds across the interface, contrasting with few hydrophobic interactions (Supplementary Table 1).

**Figure 6.**
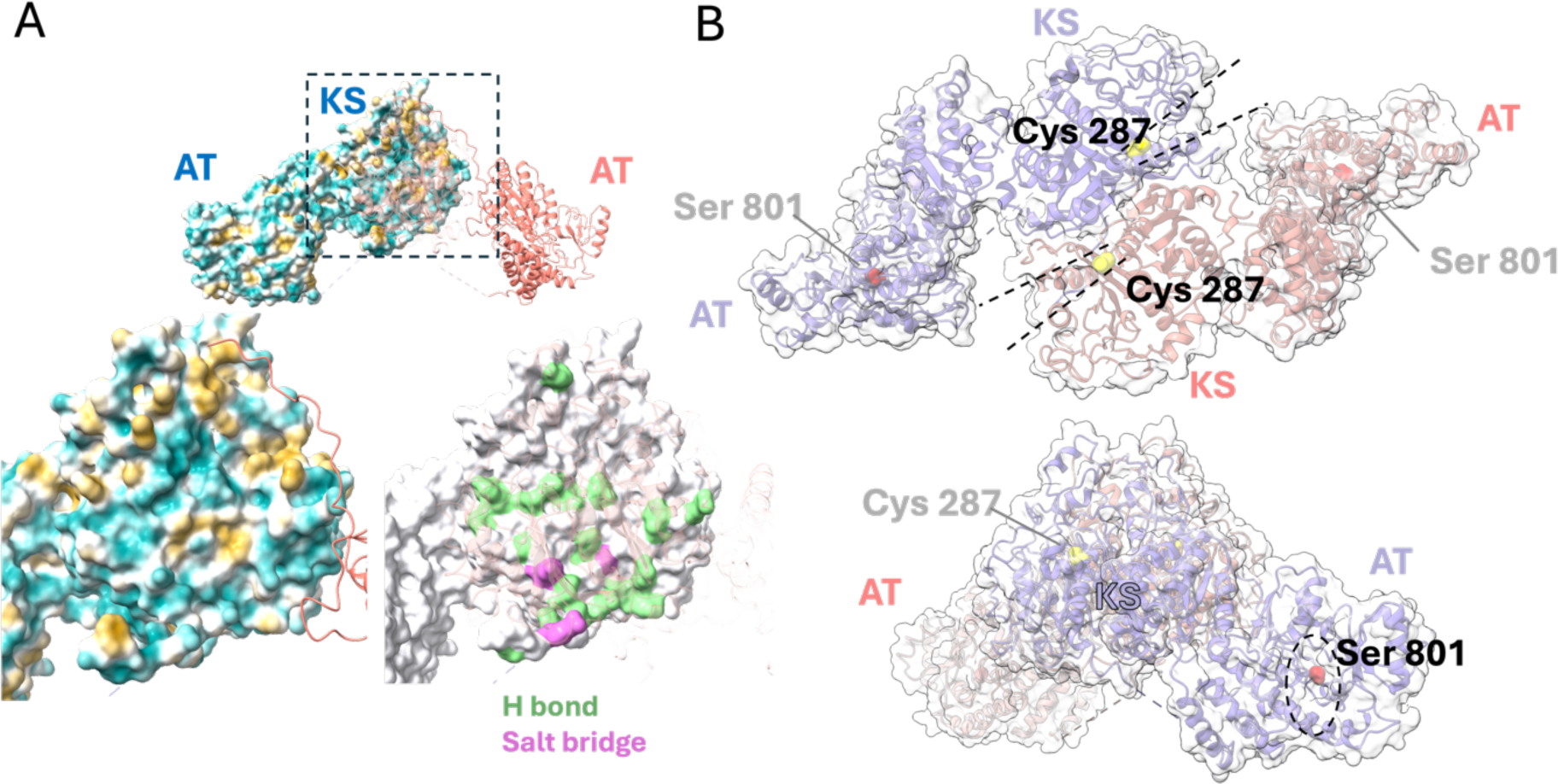
Analysis of the dimer interface and location of active sites. **A** Top: Pks13 drawn as a molecular surface (coloured according to hydrophobicity) for one monomer and a ribbon representation for other (translucent ribbon for the KS domain). Hydrophobic surfaces are coloured in gold, hydrophilic ones in light blue. Bottom left: close-up of the KS dimer interface coloured by hydrophobicity. Bottom right: close-up view of the interface highlighting residues forming H-bonds (green) and salt bridge interactions (purple) as per analysis using PISA (Supplementary Table 1). **B** Top: View of the dimer down the 2-fold rotation axis with active site cavities of the KS domains (Cys 287, yellow) indicated by dashed lines. In this orientation, the active site cavities for the AT domains are oriented away from the viewer. Bottom: active site cavity of the AT domain (Ser 801, red, dashed oval) opening towards the viewer.

The active sites of the KS and AT domains are well separated from each other and the active site cavities in the molecular surface open into diametrically opposed directions (Fig. 6B). For instance, the cavities leading to the active site cysteine (Cys 287) of the KS domain open in the direction of the AT domains of the opposing monomers, but the cavities leading to the AT domain catalytic serine (Ser 801) are opening downwards (in the orientation of Fig. 6B).

### Identification of active site residues

The cryoEM structure of Mt-Pks13 was obtained without substrates or inhibitors present in the protein solution. To identify the location of the active site, we superimposed Mt-Pks13 with ligand-bound structural homologues (Fig. 7). The KS domain closely matches the KS3 domain of the modular PKS 6-Deoxyerthronolide B synthase (DEBS), which synthesises the erythromycin core in *Saccharopolyspora erythraea* (PDB entry 2QO3). In the KS3 domain of DEBS, the inhibitor cerulenin forms a covalent bond with the catalytic cysteine of a Cys-His-His triad, whereby the two histidine residues also coordinate the inhibitor (Fig. 7A) (Tang et al., 2007). In the crystal structure of the Mt-Pks13 AT domain conjugated to carboxypalmitoyl (PDB entry 3TZZ, Bergeret et al., 2012), the alkyl chain of palmitoyl occupies a tunnel leading from the surface to the catalytic serine (Ser 801). This tunnel is occluded by residues 905-907 in our cryoEM structure (Fig. 7B, bottom right panel).

**Figure 7.**
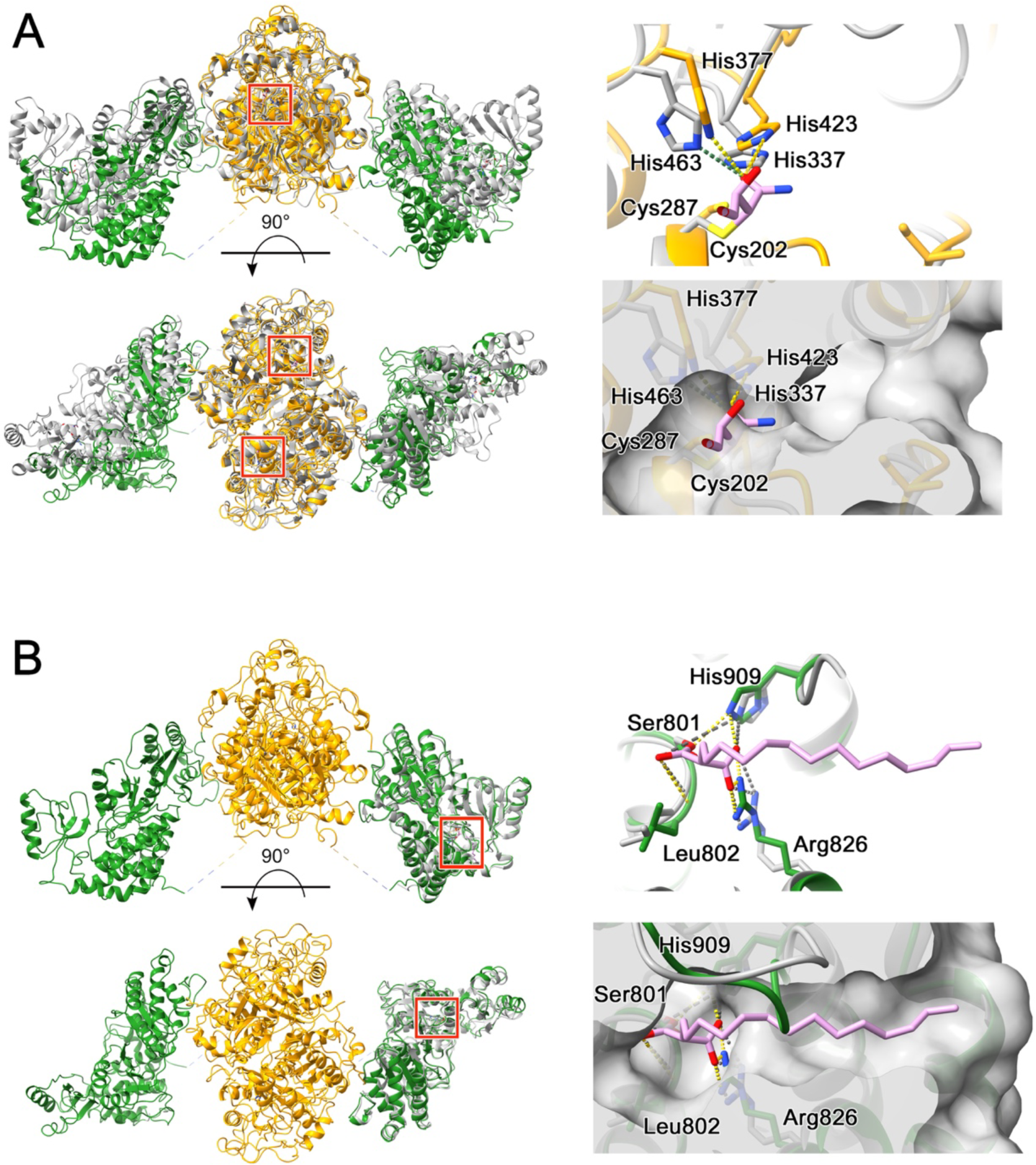
Analysis of the Mt-Pks13 KS and AT domain active site pockets. **A** Comparison with the KS3-AT3 di-domain from 6-Deoxyerthronolide B synthase (DEBS) (grey), complexed with cerulenin (pink) via the His-His-Cys triad (covalent bond with Cys202). From *Saccharopolyspora erythraea* (PDB: 2qo3) (Tang et al., 2007). **B** Superposition of the crystal structure of the carboxypalmitoylated form of the AT domain (PDB: 3TZZ, Bergeret et al., 2012) with the AT domain from the present cryoEM structures. Left panels show the alignment as a ribbon diagram with the cryoEM structure in gold/green, and the comparator structures in grey. Right, close-up views of the active site and below, surface fill of the active site pocket: in (**A**) surface fill uses the Mt-Pks13 structure, but in (**B**), loop in the cryoEM structure (residues 905-907, green) closes the active site and the pocket is from the AT crystal structure. Abbreviations KS, ketoacyl synthase (yellow) and AT, acyltransferase (green). Residues within 4Å of the compound, and suspected to be involved in coordination, are labelled with the relevant chain colour and number with a dotted line to the compound and coloured by heteroatom (red, oxygen; blue, nitrogen; yellow, sulphur). Images drawn using ChimeraX (Meng et al., 2023).

### Crossover of ACP1-KS linker sequences

In the process of constructing the structural model of Mt-Pks13, we observed that the sequences located N-terminal to the KS domain (i.e. upstream of residue Asp118) cross over to the opposite monomer within the KS domain dimer (Fig. 8A). This configuration is in stark contrast to the backbone conformation found in the structure of *M. smegmatis* Pks13 (Ms-Pks13, PDB entry 8CV1) (Kim et al., 2023).

**Figure 8.**
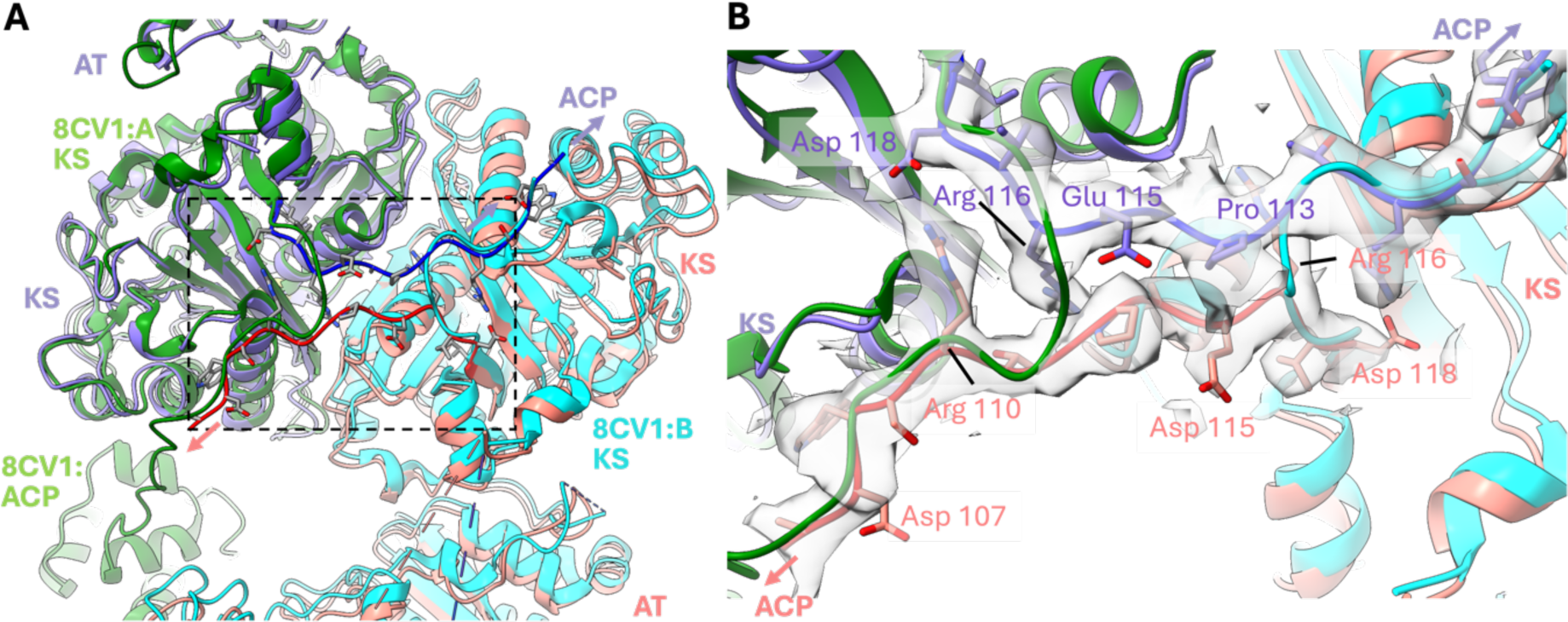
Evidence for chain crossover between KS domains. Superposition of Mt-Pks13 with the structure of M. smegmatis Pks13 (Ms-Pks13, PDB entry 8CV1, traces green, cyan,(Kim et al., 2023)). Residue numbers refer to the sequence of Mt-Pks13 **A.** View onto the N-terminal region of the KS domain with ACP-KS linker sequences crossing the domain boundary. **B**. Close-up view of the boxed region in panel A; the cryoEM density map is overlayed with the structure of Mt-Pks13 and the ribbon of Ms-Pks13 (8CV1). The C**D** traces of Ms-Pks13 show a near 180° turn near Mt-Pks13 residue Arg116. KS, ketoacylsynthase; AT, acyltransferase; ACP, acyl carrier protein.

Superimposing our model with that of Ms-Pks13, which includes the N-terminal ACP domain, we found that the backbone N-terminal to the KS domain in Ms-Pks13 makes a ∼180° turn (Fig. 8B). This turn causes the ACP domain of chain A to fold back onto the KS domain of the same subunit (Fig. 8A, trace in green). Our cryoEM map shows continuous backbone density for both subunits in the region N-terminal to residue Asp 118 that clearly indicates an extended conformation of the ACP-KS linker sequence rather than a turn. As a result, the ACP of chain A likely associates with the KS domain of the opposite subunit (and vice versa). The turn observed in Ms-Pks13 occurs near the Arg116 residue of Mt-Pks13, where density for the Arg side chain is clearly delineated and does not bridge to the backbone density of the opposite monomer. Secondly, density for large side chains in this region (e.g. Trp108, Arg110, Arg116) is well defined providing confidence that the sequence register is correct and the extended confirmation is not the result of a model building artefact.

### Additional structural features in a low resolution cryoEM map

During data processing and successive rounds of the 3D reconstruction, we obtained a low-resolution map (∼15 Å, Supplementary Fig. 1) that showed additional features compared to the eventual high-resolution map. Intriguingly, these features can be attributed to identifiable structural entities of Pks13 through modelling using AlphaFold2 (AF2) (Mirdita et al., 2022), superimposition with the KS-AT di-domain structure or direct docking into the low-resolution map.

For instance, an AF2 model for Mt-PKs13 comprising residues 1 to 600 was docked into the map of Fig. 8 by rigid-body fitting, using ChimeraX (Meng et al., 2023). This docking procedure matched the N-terminal ACP domain with an additional density blob between the AT and KS domains that is visible for only one monomer in the dimer. Similarly, superposition of our KS-AT di-domain structure with the structure of the ACP-KS-AT fragment of Ms-Pks13 (PDB entry 8CV1) (Kim et al., 2023) places the N-terminal ACP domain into the same density feature (Fig. 8A).

Secondly, the linker between the KS and AT domain (residues 551 – 595) in the AF2-generated model maps into appendix-like density features between the KS and AT domains (Fig. 9A), although the match is incomplete.

**Figure 9.**
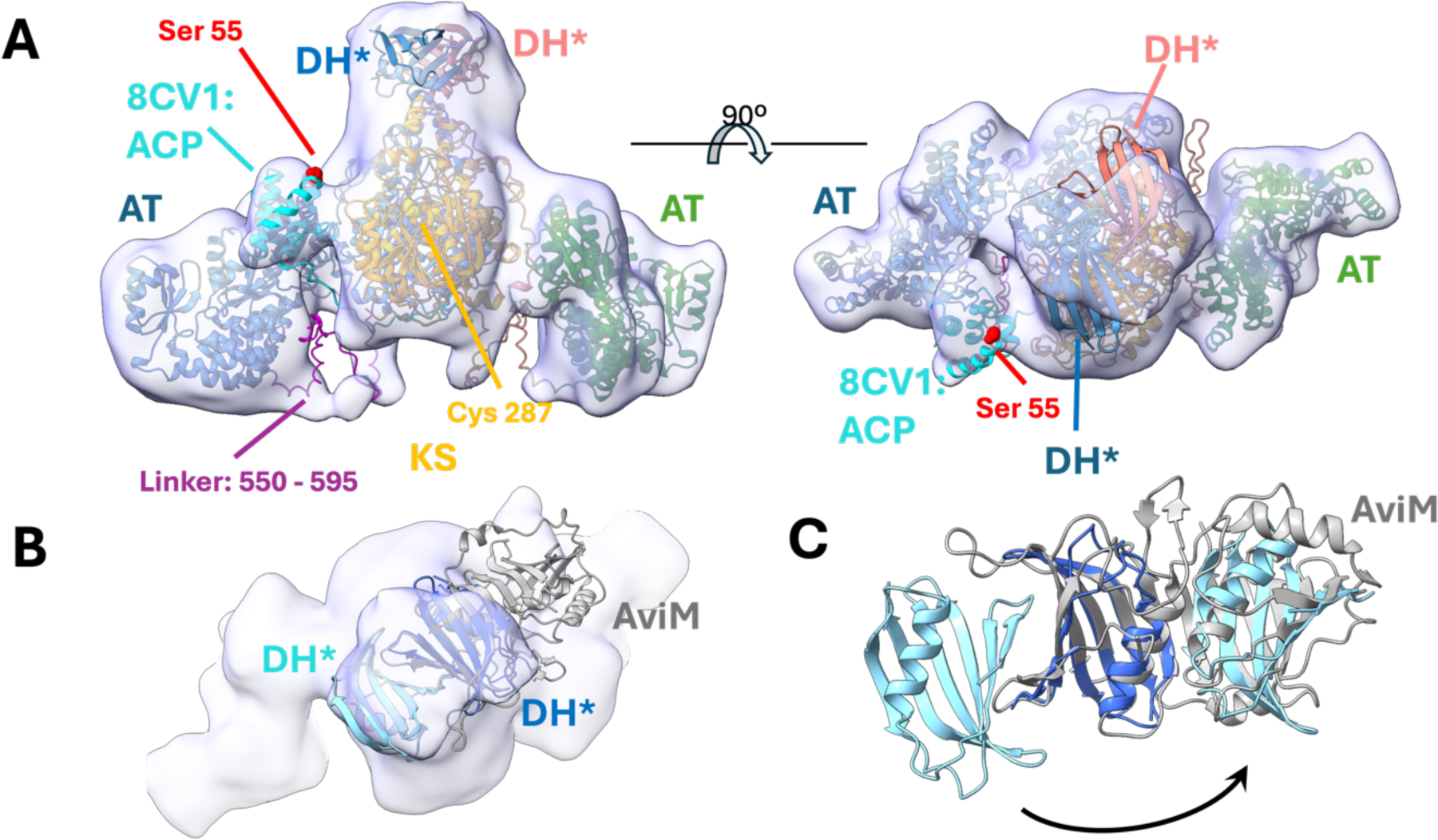
Low resolution cryoEM map of Mt-Pks13 and docking of structural models. **A** Low resolution cryoEM map (15 Å resolution) with structural features that did not carry through to the final high resolution map. Composite of structural models docked into the map using ChimeraX, (Meng et al., 2023), starting with the AlphaFold2-generated structure of the KS-AT-DH* fragment of Mt-Pks13 (labels AT, KS, DH*). The structure of Ms-Pks13 (PDB: 8CV1) comprising the N-terminal ACP, KS and AT domains was superimposed, resulting in the positioning of the ACP (cyan) into the density between the KS and AT domains. **B** Superposition of the AF2-generated model of the DH* dimer (through co-folding) with the low resolution cryoEM map and with of a DALI-derived structural neighbour, the dehydratase-like AviM product template domain from *Streptomyces viridochromogenes* (PDB: 7VWK) (Feng et al., 2022). See supplementary information for details of the DALI search. **C** Illustration of the duplication of the DH* fold within the dehydratase domain of AviM by mapping one copy of DH* onto residues 1070 – 1170 (the C-terminal half) of AviM.

Thirdly, the AF2-generated monomeric model for residues 106-1171 correctly predicted the structure of the KS-AT segment and, in addition, places a structural domain which we have designated as the DH* domain into the density on top of the KS domain. Thus, the cryoEM data capture features at low resolution that lack the definition of the ultimate high-resolution map, but that are consistent with AF2 modelling and consistent with comparison to the recently published cryoEM structure of *M. smegmatis* Pks13 (Kim et al., 2023).

### DH* domain

Given that the AF2 model for residues 106 – 1171 had predicted a hitherto unknown domain C-terminal to the AT, covering residues 1074 – 1172, we explored this AF2-predicted domain through searching for structural homologues using DALI (Holm, 2020), a search algorithm that uses distance matrix alignment to identify structural neighbours independent of primary sequence.

The DALI search (Supplementary Table 2) returned a number of high-confidence matches with dehydratase or dehydratase-like domains in PKS-like enzymes. For instance, superposition of the AF2-generated DH* domain with the dehydratase-like domain of AviM from *Streptomyces viridochromogenes* suggests that the DH* adopts a fold that represents the N-terminal ‘half’ of the dehydratase domain in AviM (Fig. 9B, C). A second copy of the DH* domain can be shoehorned on the C-terminal half of AviM (Fig. 9C), but the match is less compelling than for the N-terminal half. Given that the KS domain forms a dimer, we probed potential dimerization of the DH* domain by co-folding two copies the of the corresponding Mt-Pks13 sequence segment. Varying the sequence fragment length and using sequences from related mycobacterial species, AlphaFold2 consistently predicted the same dimeric model (Fig. 9B), which fits the density area above the KS domain dimer. Structural alignment with AviM and other structural neighbours indicates that the catalytic residues required for dehydratase activity are not conserved in the DH* domain, and thus this domain lacks an enzymatic role within Mt-Pks13.

## Discussion

The elucidation of the 3D structure of Pks13 has thus far produced only partial models of this essential enzyme, even when, as in our case, the full-length protein was flash frozen on cryoEM grids (this work and (Kim et al., 2023)). Different species or expression systems cause different preparative challenges. For instance, while purifying recombinant Mt-Pks13 from *E. coli* necessitated a reducing agent, this was not required when Ms-Pks13 was purified from its native host, *M. smegmatis* (Kim et al., 2023). Conversely in our preparation no detergent was required, unlike the procedure described for Ms-Pks13. These discrepancies could be linked to pantetheinylation of the ACP domains’ invariant serine residues, which is occurring in the native host but not when the recombinant protein is expressed in *E. coli* due to the lack of the required posttranslational modification machinery.

The relative stability of the assembly of the two core domains (KS, AT) into a dimer, which is clearly discernible among particles in 2D class averages (Supplementary Fig. 1), ostensibly does not extend to the ACP domains or the C-terminal TE domain, all of which are essential for function. Our initial assumption that an incomplete structure was obtained because Pks13 may be vulnerable to proteolytic digestion during purification is contradicted by the AUC equilibrium mass determination, which returned a monomeric species and a close match to the sequence-derived molecular mass (Fig 3). However, the solution-state analysis by AUC also established that a pH-dependent monomer-dimer equilibrium, which seems mirrored by a subpopulation of smaller particles (presumably monomers) in the 2D class averages (Supplementary Fig. 1). This finding is corroborated by a recent small angle x-ray scattering (SAXS) analysis of Pks13 (Bon et al., 2022), revealing monomers as the predominant species, with a propensity to dimer formation when ACPs are loaded with a C16 alkyl chain. Furthermore, variable orientation between the KS and AT domains was observed in the cryoEM structure of Ms-Pks13, as was distinct conformations for the N-terminal ACP in relation to the KS-AT core domains (Kim et al., 2023).

Taken together, the structural data clearly indicate that Pks13 is not a rigid macromolecular entity but incorporates considerable conformational diversity, in part mediated through the interdomain linkers and the mobility and length of the Ppant arms, a conformational freedom that is presumably mechanistically important. For instance, when the ACPs, situated on up- and down-stream of the KS-AT di-domain, are loaded with their respective acyl chains, they must successively deliver the reactive moieties of those chains to the active sites of the KS to facilitate the condensation reaction (Witkowski et al., 2002). How this occurs is readily visible for the N-terminal ACP (ACP^N^) through its position in the low-resolution map, juxtaposing the Ppant attachment site in the ACP^N^ (Ser55) to the active site cavity of the KS domain (Figs. 6, 9). Structural clues for how the C-terminal ACP (ACP^C^, residues 1234-1307) participates in catalysis are more indirect. The ACP^C^ domain is separated from the DH*, which ostensibly associates with the KS-AT di-domain core, by a linker sequence of about 60 amino acid residues (Fig. 10). The straight-line distance from the “top” of the DH* domain to the active site serine of the AT domain is approximately 105 Å, while the distance to the position of the N-terminal ACP (ACP^N^) is about 68 Å (Fig. 10). In a fully extended conformation, the DH*-ACP^C^ linker could stretch to ∼200 Å, thus is of sufficient length to allow the ACP^C^ domain to position the Ppant attachment site (Ser1266) within chemically relevant proximity of the catalytic groups in the AT or KS domains. The linker sequence between the ACP^C^ and the non-functional ACP* (upstream of the TE domain) is of similar length (Fig. 10), suggesting adequate freedom of movement or conformational leeway between the ACPs and the three catalytic domains of Pks13, even if the KS-AT di-domains were to remain conformationally constrained by dimer formation.

**Figure 10.**
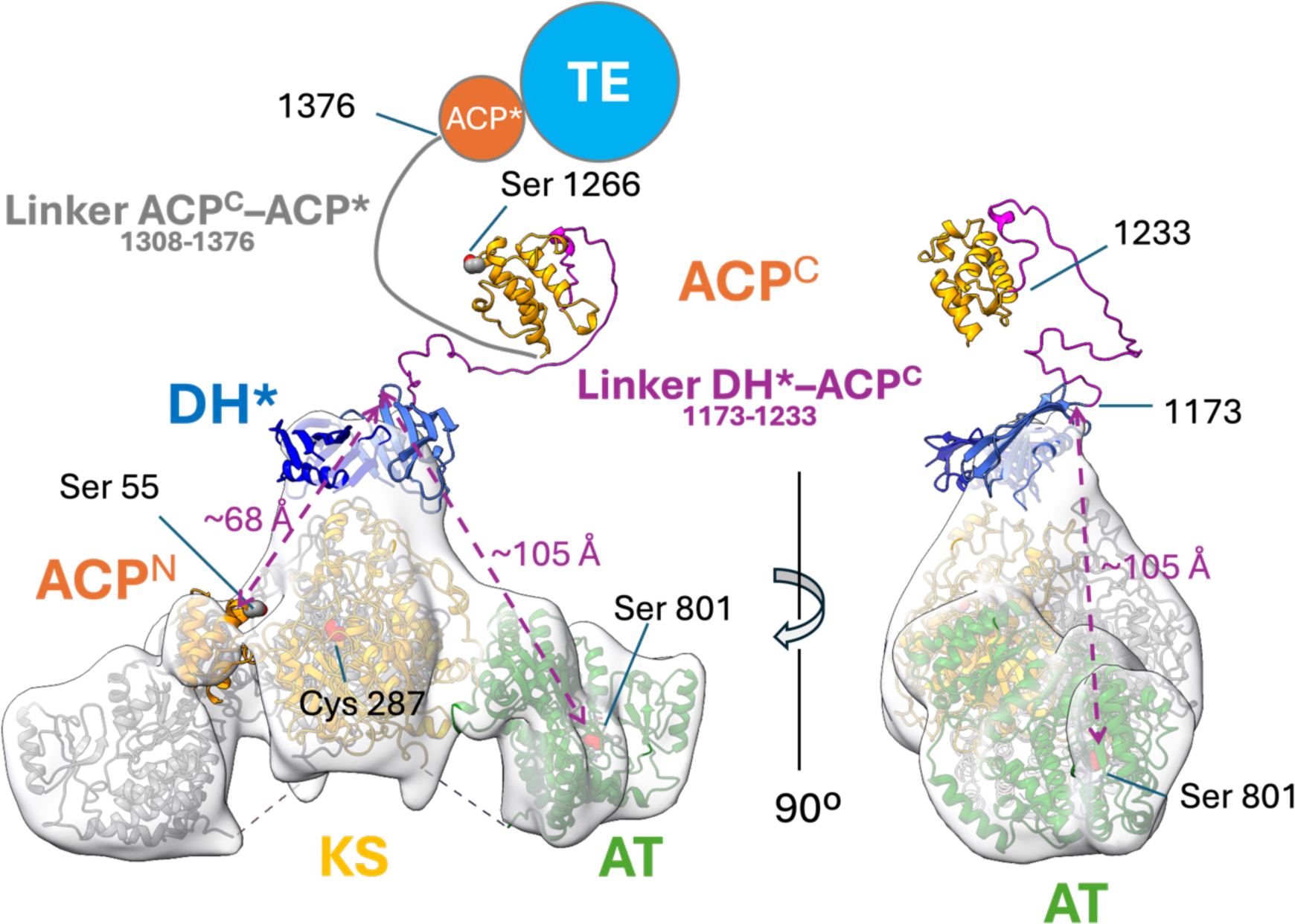
Composite model of Pks13 illustrating the putative mechanistic role of linker regions in Mt-Pks13. The final refined model of Pks13 was combined with AF2-generated models of domains, which were not observed in the high-resolution cryoEM map, and superimposed with the 15 Å intermediate map (Fig. 9A). The dashed double arrows (magenta) indicate straight-line distances between the C-terminus of the DH* domain and functionally required positions of the ACP^C^ domain at different stages of the reaction cycle. DH*, pseudo-dehydratase; ACP*, inactive ACP; KS, ketoacyl synthase; AT, acyltransferase; ACP^N,C^, N- and C-terminal ACP domains.

Furthermore, the alternative conformations of the N-terminal ACP-KS linker sequences (residues 105-118) seen in our structure (Fig. 8) *versus* that of Ms-Pks13 suggests that in the context of a dimeric assembly, the N-terminal ACP can associate with either of the KS domains in the dimer. Finally, the mono-dimer equilibrium observed in our AUC experiment and corroborated by the SAXS data (Bon et a, 2022) raise the question whether enzymatic activity is mediated by either form, or mechanistically tied to the more common dimeric assembly observed in other PKS-type enzymes (Robbins et al., 2016). For instance, the PKS CalA3 has a similar dimeric architecture as Pks13, with a sequence of KS-AT-DH-KR*-KR domains, where KR stands for ketoreductase activity. CalA3’s DH domain is located above the KS-AT di-domain and participates in dimer formation, in analogy to the putative DH* domain in Mt-Pks13 (Wang et al., 2023). A second fascinating parallel is the presence of a catalytically dysfunctional KR (referred to as KR*) in CalA3, analogous to the DH* and inactive ACP* (residues 1376-1447) in Mt-Pks13.

Mt-Pks13 has emerged as a prominent target in phenotypic screens identifying novel anti-tubercular inhibitors, an occurrence closely linked to the pathogen’s need for an intact mycolic acid layer to survive host cell stresses (Batt, Burke, et al., 2020). Potent anti-mycobacterial compounds directed at Pks13 include benzofurans, coumestan, oxadiazoles and β-lactones which bind to the C-terminal TE domain (Aggarwal et al., 2017; Green et al., 2023; Lehmann et al., 2018; W. Zhang et al., 2018), while thiophene compounds prevent meromycolyl chain attachment to the N-terminal ACP (Wilson et al., 2013). These examples illustrate a broad spectrum of options to interfere with Pks13 function, from targeting inhibitors to one of the three catalytic sites, to preventing substrate delivery, perhaps even the option to increase specificity through multi-mode inhibitors.

In summary, our study of Mt-Pks13 revealed the molecular structure of the KS-AT di-domain, evidence for conformational flexibility and a propensity to switch between monomeric and dimeric assembly states in response to pH. The availability of diverse experimental structures of this promising target for antimycobacterial therapy gives options for computer-aided drug discovery or design, and in this way can contribute to meeting the WHO Sustainable Development Goals (WHO, 2016) in relation to the global TB epidemic.

## Materials and Methods

### Overexpression of Pks13 in *E. coli*

The pET23b-Pks13 plasmid vector had been previously constructed using the primers 5’GATCGATC<u>CATATG</u>GCTGACGTAGCGGATTC-3’ and 5’-GACTGATC<u>AAGCTT</u>CTGCTTGCCTACCTCACTTG-3’, cloning into the vector using the underlined NdeI and HindIII sites (Brown, 2004). The recombinant protein, Mt-Pks13 was expressed in *E. coli* C41 (DE3) cells cultured in 1 L Terrific broth (Difco). Cultures were grown at 37°C to mid-log phase (A_600_=0.4-0.6) and induced overnight at 18°C with 0.5 mM isopropyl-β-D-1-thiogalactopyranoside. Cells were harvested by centrifugation (6,000 ×g, 10 min, 4°C), washed in Phosphate Buffered Saline and stored at −20°C.

### Purification of Pks13 from cell extracts

The cell pellets were resuspended in 30 ml buffer A (20 mM NaHPO_4_, 500 mM NaCl, pH 7.9, 2.5 mM β-mercaptoethanol (βME)), supplemented with 1 mM phenylmethylsulfonyl fluoride (PMSF), and a cOmplete, an EDTA-free protease inhibitor cocktail tablet (Roche). Cells were lysed by sonication and the insoluble material pelleted by centrifugation (48,000 ×g, 45 min, 4°C). The supernatant was syringe filtered through a 0.45 µm filter before purifying recombinant Mt-Pks13 through a 5 mL HiTrap Ni-NTA column (Cytiva). The column was pre-treated with buffer A plus 500 mM imidazole and equilibrated with buffer A prior to loading the lysate. Mt-Pks13 was eluted with a gradient of imidazole in buffer A. Fractions were collected and analysed by SDS-PAGE.

Elution fractions containing Mt-Pks13 were pooled and dialysed overnight in buffer B (20 mM Tris-HCl pH 7.9, 50 mM NaCl, 2.5 mM βME). The protein was then loaded onto a 5 ml HiTrap Q-sepharose HP anion exchange chromatography (QHP) column (Cytiva), pre-equilibrated with buffer B. Mt-Pks13 was eluted with increasing concentrations of NaCl in buffer B up to 1 M NaCl. Fractions were collected and analysed by SDS-PAGE.

Fractions containing Pks13 were concentrated using a 100K MWCO centrifugal filter unit (Merck Millipore) and further purified by size exclusion chromatography using either a HiLoad Superdex 200 26/60 PG or Superdex 200 increase 10/300 (Cytiva) column in buffer B with 150 mM NaCl.

### Sedimentation equilibrium analytical ultracentrifugation

Mt-Pks13 QHP elution fraction was diluted to an absorbance at 280 nm (A_280_) of 0.75, 0.5 and 0.25 in buffer B with 500 mM NaCl. The samples were run in a 6-well cuvette, with corresponding buffer at 4°C and speeds of 10,000, 12,000, 14,000 rpm using a Beckman Coulter ProteomeLab XL-1 analytical centrifuge. The absorbance data was then initially analysed using SEDFIT (Schuck, 2000) to set the bottom and meniscus for each sample well. This data was then transferred to SEDPHAT (Schuck et al., 2023) and analysed and fitted globally across all speeds at each concentration, using the species analysis model to estimate the molecular weight and statistical analysis performed using Monte-Carlo error analysis (over 1000 iterations, confidence level of 0.68). The buffer parameters for analysis were calculated using UltraScan (Scott et al., 2005) (partial specific volume of 0.7324 mL/g, buffer density of 1.01924 g/cm^3^, and buffer viscosity of 1.04992 cP).

### Sedimentation velocity analytical ultracentrifugation

Purified Mt-Pks13 was dialysed into a triple buffer containing 20 mM each of Sodium Acetate: Tris-HCl: N-Cyclohexyl-2-aminoethanesulfonic acid (CHES), 150 mM NaCl, and 2.5 mM βME, with buffers at varied pH values between pH 3.5-10. Each protein sample was analysed by sedimentation velocity AUC using a Beckman Coulter ProteomeLab XL-1 analytical centrifuge, at either 4°C or 20°C, 30,000 rpm. Data was then analysed using SEDFIT using the continuous c(S) distribution model (Schuck, 2000; Schuck et al., 2002). Buffer parameters were calculated using UltraScan (Scott et al., 2005) (partial specific volume of 0.7324 mL/g, buffer density of 1.00608 g/cm^3^, and buffer viscosity of 1.02009 cP).

### Cryo-EM data acquisition and processing

A volume of 3 µL of purified Pks13 at 0.9 mg/ml was loaded onto a glow discharged (35 mA, 60 seconds) R1.2/1.3 Au grid with holey carbon foil (Quantifoil). This was blotted for 3 sec at 100% humidity and plunge frozen in liquid ethane using a Vitrobot Mark IV system (ThermoFisher).

Super-resolution micrographs were recorded using a FEI Titan Krios G3 at 300 kV with a Gatan K3 detector (Midlands Regional Cryo-EM Facility), using EPU software. Data was collected at a nominal magnification of 81,000 ×, a 1.086 Å pixel size and a dose rate on the specimen of 16.5 e^−^/pix/sec. Images were collected in 43 fractions with a 3 s exposure time. A total of 1,412 movies were collected over a defocus range of –2.7 to –1.5 µm.

All data processing of the cryo-EM micrographs was completed using Relion 3.1 (Scheres, 2012). Micrographs were motion corrected using Relion’s implementation of MotionCor2 (Zheng et al., 2017). CTF estimation was performed with Gctf (Zhang, 2016), and micrographs with poor CTF estimates were excluded manually. Topaz picker version 0.2.4 (Bepler et al., 2019) was used to automatically pick 310,055 particles. Extracted particles were subjected to multiple rounds of 2D classification using a mask diameter of 190 Å and the final classes were used to form a 3D initial reconstruction model in Relion. 3D classification was performed using the initial model as a reference and the best class which resembled a dimer (containing 34,757 particles) was selected for further refinement (see Supplementary Fig. 1). At this stage, a 15 Å resolution 3D model was identified to contain density which was attributed to the N-terminal ACP domain, however this was lost during subsequent refinement and processing steps. Next the particles were used for an initial 3D refinement step, with a 210 Å mask diameter. A solvent mask was then created using a 15 Å low-pass filtered map to determine the initial binarization threshold in Chimera (Pettersen et al., 2021), and the unfiltered half-maps subjected to post-processing using this mask. This loop was then repeated using the mask at 3D refinement. CTF refinement was then completed on the refined particles and the loop repeated.

Bayesian polishing was added to further improve map quality, re-extracting the particles at 1.086 Å/pix (Scheres, 2012; Zivanov et al., 2018). This pipeline of 3D and CTF refinement followed by polishing was repeated multiple times until no improvement in resolution was seen.

### Model building

The di-domain KS-AT core were built de novo, guided by the KS-AT structure of 6-deoxyerthronolide B synthase (PDB 2H4G) and the crystal structure of Pks13 AT domain (PDB 3TZY). Model building was performed using Coot (Emsley et al., 2010) with subsequent real space refinement using the Phenix package (Liebschner et al., 2019).

### Database deposition

Coordinates of the structure of the Mtb-Pks13 KS-AT di-domain have been deposited in the Protein Data Bank (PDB, wwPDB.org) under accession code 9F48. The corresponding cryoEM density map has been deposited in the EMDB under accession code EMD-50185.

## Acknowledgments

Funding has been awarded as follows: Personal Research Chair from Mr. James Bardrick, G.S.B. Medical Research Council MR/S000542/1, G.S.B. Medical Research Council MR/R001154/1, G.S.B. Sir Charles Hercus Health Research Fellowship awarded through the Health Research Council of New Zealand, G.B.

## Supplementary Information

Supplementary figures and tables are available for this manuscript.

