## Supplemental for "CryoEM structure of the di-domain core of *Mycobacterium tuberculosis* polyketide synthase 13, essential for mycobacterial mycolic acid synthesis"

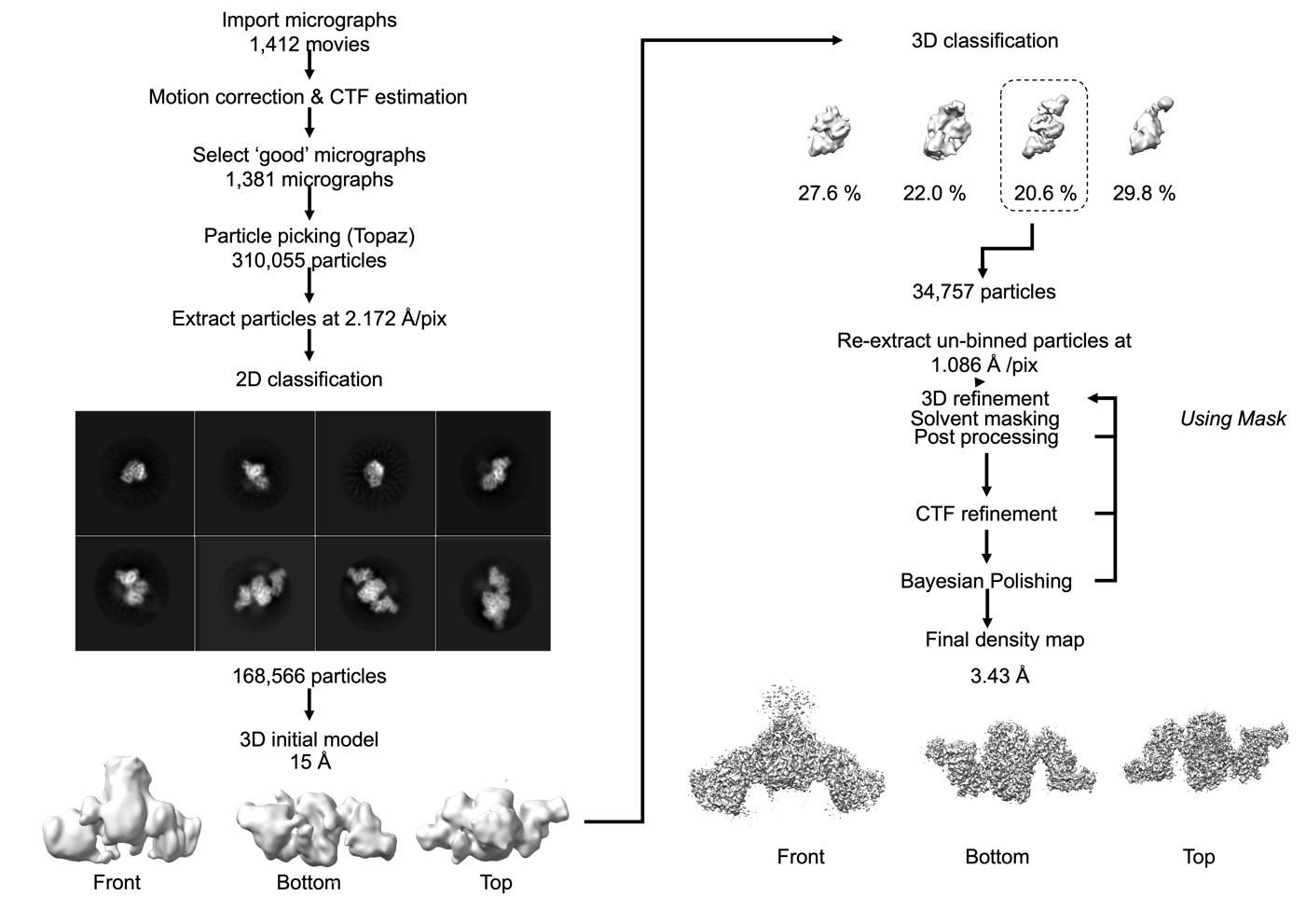

**Supplementary Figure 1. CryoEM workflow used to derive the electron density map for Mt-Pks13.**

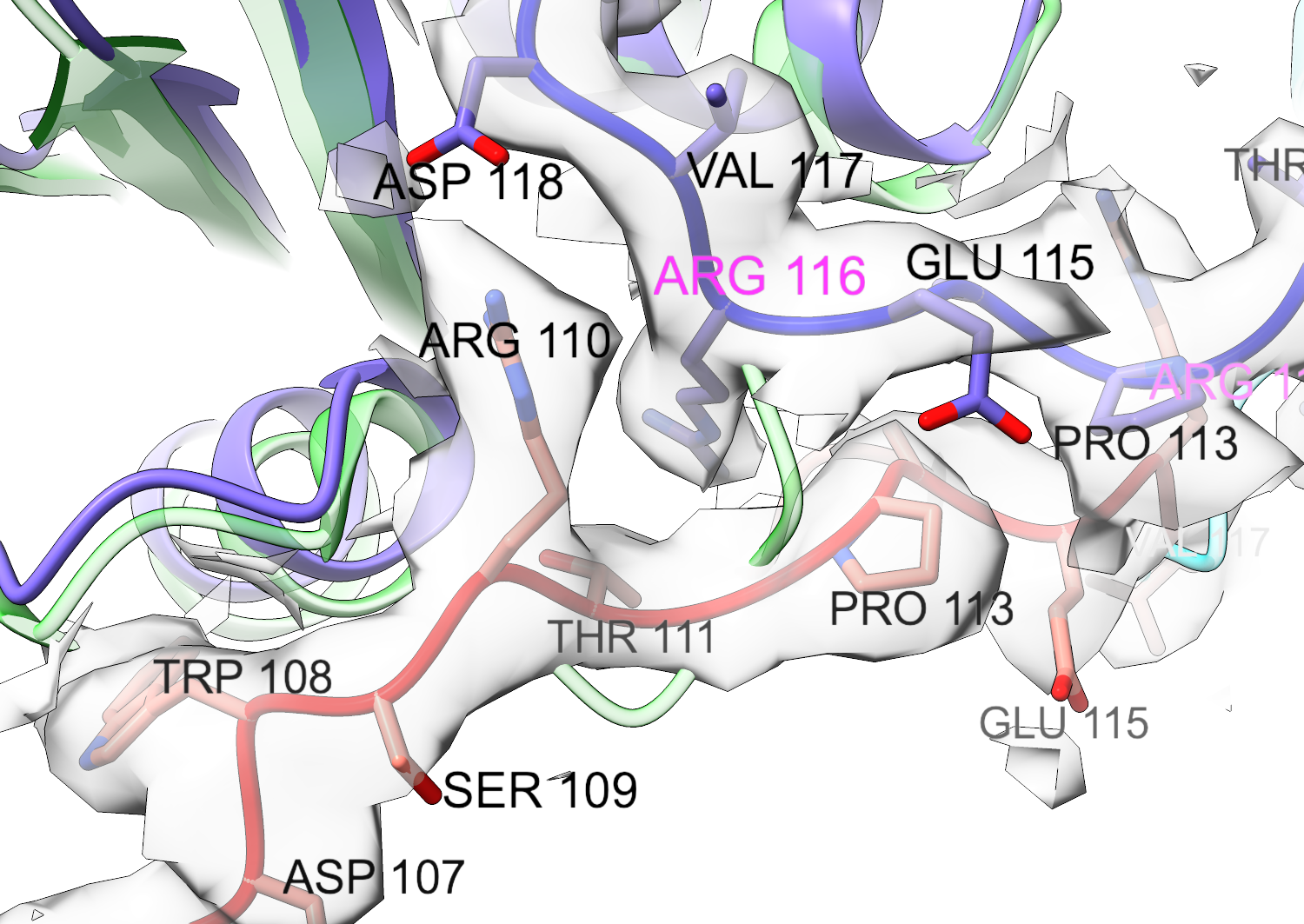

**Supplementary Figure 2. Density map for Mt-Pks13 around residue Arg116.** Ribbon in green is the Cα trace of Ms-Pks13 (PDB entry 8CV1, (Kim et al., 2023)). Traces in blue and red represent chains A and B, respectively, of Mt-Pks13. Residue numbers refer to the sequence of Mt-Pks13.

**Supplementary Table 1. Analysis of the KS-KS dimer interface using PISA (ref).** Listing of H-bond and salt bridge interactions detected by the software.

| **H-bond interactions** | | | |
| --- | --- | --- | --- |
| **##** | KS domain (chain A) | Dist. [Å] | KS domain (chain B) |
| 1 | A:ARG 116[ NH1] | 2.40 | B:GLY 112[ O  ] |
| 2 | A:ALA 114[ N  ] | 3.51 | B:ALA 114[ O  ] |
| 3 | A:ARG 110[ NH2] | 2.73 | B:ASP 118[ OD2] |
| 4 | A:ARG 387[ NH2] | 2.65 | B:PHE 273[ O  ] |
| 5 | A:ARG 387[ N  ] | 2.74 | B:ASP 275[ OD1] |
| 6 | A:HIS 384[ N  ] | 3.45 | B:GLY 278[ O  ] |
| 7 | A:ASP 284[ N  ] | 3.43 | B:THR 282[ O  ] |
| 8 | A:THR 282[ N  ] | 3.61 | B:ASP 284[ O  ] |
| 9 | A:SER 264[ N  ] | 3.07 | B:ASP 284[ OD2] |
| 10 | A:ARG 116[ NH2] | 3.53 | B:ASN 303[ O  ] |
| 11 | A:ARG 405[ NH2] | 2.76 | B:GLU 305[ O  ] |
| 12 | A:ARG 110[ NH1] | 3.12 | B:ALA 378[ O  ] |
| 13 | A:THR 111[ N  ] | 3.21 | B:LYS 408[ O  ] |
| 14 | A:THR 111[ OG1] | 2.93 | B:LYS 408[ O  ] |
| 15 | A:ARG 110[ NH1] | 2.85 | B:ASP 409[ O  ] |
| 16 | A:ASP 107[ O  ] | 3.38 | B:LYS 408[ NZ ] |
| 17 | A:GLY 112[ O  ] | 3.22 | B:ARG 116[ NE ] |
| 18 | A:ALA 114[ O  ] | 3.15 | B:ALA 114[ N  ] |
| 19 | A:ASP 118[ OD1] | 3.09 | B:ARG 110[ NH2] |
| 20 | A:PRO 250[ O  ] | 2.77 | B:ARG 169[ NH1] |
| 21 | A:ASN 268[ O  ] | 3.75 | B:HIS 384[ NE2] |
| 22 | A:TYR 272[ O  ] | 3.59 | B:ARG 387[ NE ] |
| 23 | A:PHE 273[ O  ] | 2.56 | B:ARG 387[ NH2] |
| 24 | A:ASP 275[ OD1] | 2.67 | B:ARG 387[ N  ] |
| 25 | A:GLY 278[ O  ] | 3.26 | B:HIS 384[ N  ] |
| 26 | A:THR 282[ O  ] | 3.48 | B:ASP 284[ N  ] |
| 27 | A:ASP 284[ O  ] | 3.48 | B:THR 282[ N  ] |
| 28 | A:ASP 284[ OD2] | 2.20 | B:SER 263[ OG ] |
| 29 | A:GLN 296[ OE1] | 3.11 | B:GLN 296[ NE2] |
| 30 | A:ASN 303[ O  ] | 2.98 | B:ARG 116[ NH2] |
| 31 | A:GLU 305[ O  ] | 3.87 | B:ARG 405[ NH2] |
| 32 | A:GLU 305[ OE2] | 3.29 | B:HIS 295[ NE2] |
| 33 | A:ALA 378[ O  ] | 3.12 | B:ARG 110[ NH1] |
| 34 | A:LYS 408[ O  ] | 3.37 | B:THR 111[ OG1] |
| 35 | A:LYS 408[ O  ] | 3.40 | B:THR 111[ N  ] |
| 36 | A:ASP 409[ O  ] | 3.45 | B:ARG 110[ NH1] |
| 37 | A:ASP 409[ OD1] | 3.86 | B:THR 111[ OG1] |
| **Saltbridge interactions** | | | |
| **##** | KS domain (chain A) | Dist. [Å] | KS domain (chain B) |
| 1 | A:ARG 110[ NH2] | 2.73 | B:ASP 118[ OD2] |
| 2 | A:HIS 295[ NE2] | 3.98 | B:GLU 305[ OE2] |
| 3 | A:ASP 118[ OD1] | 3.09 | B:ARG 110[ NH2] |
| 4 | A:ASP 118[ OD2] | 3.93 | B:ARG 110[ NH1] |
| 5 | A:ASP 118[ OD2] | 3.36 | B:ARG 110[ NH2] |
| 6 | A:GLU 305[ OE1] | 3.69 | B:HIS 295[ NE2] |
| 7 | A:GLU 305[ OE2] | 3.29 | B:HIS 295[ NE2] |

**Supplementary Table 2.** Identifying structural neighbours of the DH* domain using distance matrix alignment as implemented at the DALI server (Holm, 2020). The query structure was the AlphaFold2-generated structural model of the DH* domain of Mt-Pks13 (residues 1072-1171).

| **No.** | **Chain** | **Z** | **rmsd** | **lali** | **nres** | **%id** | **Description** |
| --- | --- | --- | --- | --- | --- | --- | --- |
| 1 | 7zsk-A | 11.7 | 1.9 | 85 | 1482 | 5 | PUTATIVE POLYKETIDE SYNTHASE; |
| 2 | 5il5-B | 11 | 2 | 84 | 257 | 14 | MLND; |
| 3 | 6b2v-A | 10.9 | 1.8 | 82 | 267 | 16 | SORB; |
| 4 | 7vwk-A | 10.8 | 1.9 | 84 | 260 | 14 | POLYKETIDE SYNTHASE; |
| 5 | 3cjy-A | 10.8 | 1.8 | 81 | 253 | 15 | PUTATIVE THIOESTERASE; |
| 6 | 5bp3-A | 10.5 | 1.6 | 83 | 286 | 20 | MYCOCEROSIC ACID SYNTHASE-LIKE POLYKETIDE SYNTHASE; |
| 7 | 3nwz-A | 10.2 | 1.9 | 83 | 155 | 14 | BH2602 PROTEIN; |
| 8 | 3gek-A | 10.2 | 1.8 | 85 | 132 | 12 | PUTATIVE THIOESTERASE YHDA; |
| 9 | 5kku-B | 10.1 | 1.8 | 79 | 289 | 13 | POLYKETIDE SYNTHASE TYPE I; |
| 10 | 4a12-B | 10.1 | 2 | 83 | 186 | 7 | TRANSCRIPTION FACTOR FAPR; |
| 11 | 7cpx-A | 10 | 2.4 | 89 | 2262 | 10 | LOVASTATIN NONAKETIDE SYNTHASE, POLYKETIDE SYNTHA |
| 12 | 3esi-A | 9.9 | 2.2 | 85 | 124 | 6 | UNCHARACTERIZED PROTEIN; |
| 13 | 2prx-A | 9.8 | 1.9 | 81 | 114 | 10 | THIOESTERASE SUPERFAMILY PROTEIN; |
| 14 | 2gvh-B | 9.7 | 2.7 | 87 | 250 | 6 | AGR_L_2016P; |
| 15 | 3kg7-C | 9.6 | 2 | 85 | 287 | 16 | CURH; |
| 16 | 3f1t-B | 9.5 | 2.1 | 83 | 137 | 5 | UNCHARACTERIZED PROTEIN Q9I3C8_PSEAE; |
| 17 | 3oml-A | 9.4 | 2.4 | 79 | 532 | 10 | PEROXISOMAL MULTIFUNCTIONAL ENZYME TYPE 2, CG3415 |
| 18 | 1tbu-B | 9.4 | 1.8 | 76 | 98 | 7 | PEROXISOMAL ACYL-COENZYME A THIOESTER HYDROLASE |
| 19 | 4w78-F | 9.3 | 1.9 | 77 | 127 | 6 | HYDRATASE CHSH1; |
| 20 | 3e29-B | 9.3 | 2.1 | 85 | 135 | 16 | UNCHARACTERIZED PROTEIN Q7WE92_BORBR; |
| 21 | 1c8u-A | 9.3 | 2.1 | 81 | 285 | 7 | ACYL-COA THIOESTERASE II; |
| 22 | 3bbj-A | 9.2 | 2 | 78 | 268 | 5 | PUTATIVE THIOESTERASE II; |
| 23 | 3f5o-A | 9.1 | 2.2 | 86 | 138 | 15 | THIOESTERASE SUPERFAMILY MEMBER 2; |
| 24 | 2fs2-B | 9.1 | 2.3 | 86 | 138 | 14 | PHENYLACETIC ACID DEGRADATION PROTEIN PAAI; |
| 25 | 3e8p-A | 9.1 | 2.1 | 82 | 153 | 5 | UNCHARACTERIZED PROTEIN; |
| 26 | 1vi8-B | 9 | 1.9 | 83 | 146 | 5 | HYPOTHETICAL PROTEIN YDII; |
| 27 | 5e1v-B | 8.8 | 2.5 | 82 | 274 | 7 | POLYKETIDE SYNTHASE PKSL; |
| 28 | 3dkz-A | 8.8 | 2.1 | 81 | 125 | 10 | THIOESTERASE SUPERFAMILY PROTEIN; |
| 29 | 2qwz-A | 8.8 | 2.1 | 86 | 144 | 8 | PHENYLACETIC ACID DEGRADATION-RELATED PROTEIN; |
| 30 | 3e1e-C | 8.8 | 2.3 | 86 | 141 | 10 | THIOESTERASE FAMILY PROTEIN; |
